# Evidence of Gait Impairment After mTBI in an Adolescent Pig Model

**DOI:** 10.1101/2025.03.31.646330

**Authors:** Neha Vizzeswarapu, Sloan Kanat, Tamara Reid Bush, Galit Pelled

**Author notes:** Corresponding author: Galit Pelled, PhD. 775 Woodlot Dr., East Lansing, MI, USA.

## Abstract

This study investigates the effects of mild traumatic brain injury (mTBI) on gait in Yucatan minipigs, a relevant animal model for assessing balance impairments after TBI. Retro-reflective markers were placed on different anatomical locations on the pig, and eight motion capture cameras were set up along the sides of the pig walking path to capture natural gait data. Pigs were tested both before and after induced head injury, allowing for direct comparison of gait alterations. Our findings reveal significant impairments, including reduced shoulder flexibility, altered head movement, and compensatory stabilization strategies, such as increased vertical nodding. These observations suggest that, like human mTBI patients, pigs exhibit balance deficits and modified movement patterns. The results demonstrate the utility of the adolescent pig model in studying mTBI-related balance disturbances, providing valuable insights into its neurological and biomechanical consequences. This research lays the groundwork for future studies focused on developing diagnostic and therapeutic strategies for mTBI-induced gait impairments.

## Introduction

Traumatic brain injury (TBI) is a major cause of death and disability in the United States. There were approximately 214,110 TBI-related hospitalizations in 2020 and 69,473 TBI-related deaths in 2021 in both adults and children. Unfortunately, TBI-related visits and deaths have increased from 2006 to 2014 (CDC, 2024). The most common mechanisms of injury are accidental falls, unintentionally struck by an object, motor vehicle accident, assault, and intentional self-harm. About 80% of these causes of TBI are mild while the other 20% are equally split between moderate and severe (1). Traumatic brain injuries disrupt the normal cellular function within the brain causing changes in the way neurons act and the wiring of the brain. There is an inherent risk in many sports such as football, soccer, ice hockey, wrestling, softball, and equestrian for repetitive head trauma for athletes. Mild TBI (mTBI), which accounts for the majority of all traumatic brain injuries, is often associated with symptoms like headache, dizziness, and balance disturbances.

Hospitalization for TBI is most observed in adolescents. The main causes of pediatric TBI include sports injuries, falls, and motor vehicle collisions. Children aged 0 to 4 and 15 to 19 display the highest TBI incidence rates (2). The recovery process in children is more variable than in adults. Children who sustain a TBI, demonstrate a slower rate of recovery. They experience difficulties focusing, recalling memories and psychological issues as these children are at greater risk of developing psychiatric disorders after injury (3- 8). Clinical studies show that children and youth who experienced mTBI often face motor deficits in fine motor control such as balance,gait and strength which can persist beyond the first year postinjury (9, 10).

Mild traumatic brain injury (mTBI) can result in a variety of lingering symptoms, with balance impairments being particularly common (11, 12). Patients with mild TBI often report difficulty maintaining equilibrium, which can manifest as dizziness, unsteadiness, or a sensation of “floating.” These balance disturbances are thought to arise from disruptions in the vestibular system, as well as altered sensory integration processes that involve visual, proprioceptive, and auditory inputs (13). Even though mTBI typically does not result in overt structural damage to the brain, it can impair the brain's ability to coordinate and process sensory feedback, leading to difficulties with postural control. As a result, these balance deficits can interfere with daily activities, contributing to an increased risk of falls and diminished quality of life. Kinematics and gait analysis provide valuable tools for objectively identifying and measuring balance impairments in individuals with mTBI. By utilizing motion capture systems and force platforms, kinematic analysis can track and quantify joint angles, stride length, and body posture during movement, allowing for precise identification of deviations in normal gait patterns. These deviations, such as altered step length, asymmetry in limb movements, or irregularities in trunk stability, are often indicative of balance disturbances following mild TBI. Gait analysis further enables the assessment of temporal and spatial parameters, such as walking speed and cadence, which may be affected by the injury. When combined, these analyses offer a comprehensive and objective approach to measuring balance deficits, providing clinicians with critical data for diagnosis, monitoring recovery, and evaluating the efficacy of rehabilitation interventions.

Usage of animal models can help us understand the effects of TBI such as kinematics and cognitive skills (14-18). Pigs are a good model for translational research as there are notable similarities between pig and human neuroanatomy, physiology, and behavior. The pig brain shares many structural and functional similarities with a human brain which can help translate TBI injury from a pig to a human child (19-28). For example, usage of pigs to study gait pre- and post- ischemic stroke as severity of neural injury for stroke can be assessed through changes in gait patterns (29). These changes can be compared to human stroke patients as pigs are considered to be clinically relevant animal models for humans. Given that gait and motor changes are common symptoms of TBI, pigs could also an effective model for studying these impacts in relation to human injury

The goal of this work was to use motion capture cameras to gather 3D kinematic data to analyze hoof changes, and changes in relative rotation of body segments in adolescent Yucatan minipigs before and after mTBI.

## Materials and Methods

All procedures were approved by the Institutional Animal Care and Use Committee at Michigan State University. Two female 13-weeks old Yucatan pigs were involved in this study: One received an mTBI and the other was Sham-control. Motion capture was recorded at 13-weeks old before mTBI was induced, and at 17-weeks old, a week after mTBI was induced. These pigs were housed together throughout the study. Prior to surgery, pigs were administered midazolam (0.2-0.4 mg/kg) and butorphanol (0.2-0.4 mg/kg) intramuscularly. Pigs were mask induced, intubated and inhalant anesthesia was used to maintain a surgical plane of general anesthesia. Following impact, the periosteum and skin were closed in separate layers. Inhalant anesthesia was discontinued, and the pigs were extubated after they showed signs of recovering from anesthesia. Sham-control pigs were anesthetized and intubated for the same duration of the injury protocol and underwent the same skin and periosteal incision and closure and received the same perioperative and operative medications. The closed-head injury was induced in anesthetized pigs using an electromagnetic driven impactor, as was described in detail in Islam et al. (21). Impact was delivered over the frontal cortex. This mTBI resembles an injury with a clinically relevant magnitude of head injuries reported in youth sports (30-32).

Eight motion capture cameras (Qualisys, Gothenburg, Sweden) were set up along the sides of the pig walking path ass shown in **Figure 1**. A commercial floor mat was placed on the floor of the walking path to prevent slipping of the pig hooves. Four cameras were placed at a higher height and four were placed at a lower height to optimize viewing of the pig in the walking path. Retro-reflective markers were placed on anatomical locations on the pig. Eight individual markers were placed laterally on each knee joint and on the outside of each hoof. Three marker pods, each consisting of four retro-reflective markers fixed to a flat surface, referred to as rigid bodies, were secured to the pigs in three locations. The rigid bodies were secured on top of the pigs’ head, and on the back between the shoulders and the hips **(Fig. 1B**). The markers and rigid bodies were secured to the pig skin using Coban tape. The hoof markers were fixed to the hooves using double sided tape. Each pig was wearing a harness, but the harness was only used if the pig began to leave the designated path and needed to be redirected. All pigs were familiarized with the testing space before the mTBI procedure. The pigs were trained to follow a trainer through the pig walking path and were rewarded with treats. During testing, the pigs walked back and forth through the walking path until adequate passes of the pig through the calibrated space were collected. The cameras recorded in continuous capture at 100Hz. The 3D location (x, y and z coordinates) of each marker was exported and analyzed via MATLAB software.

**Figure 1.**
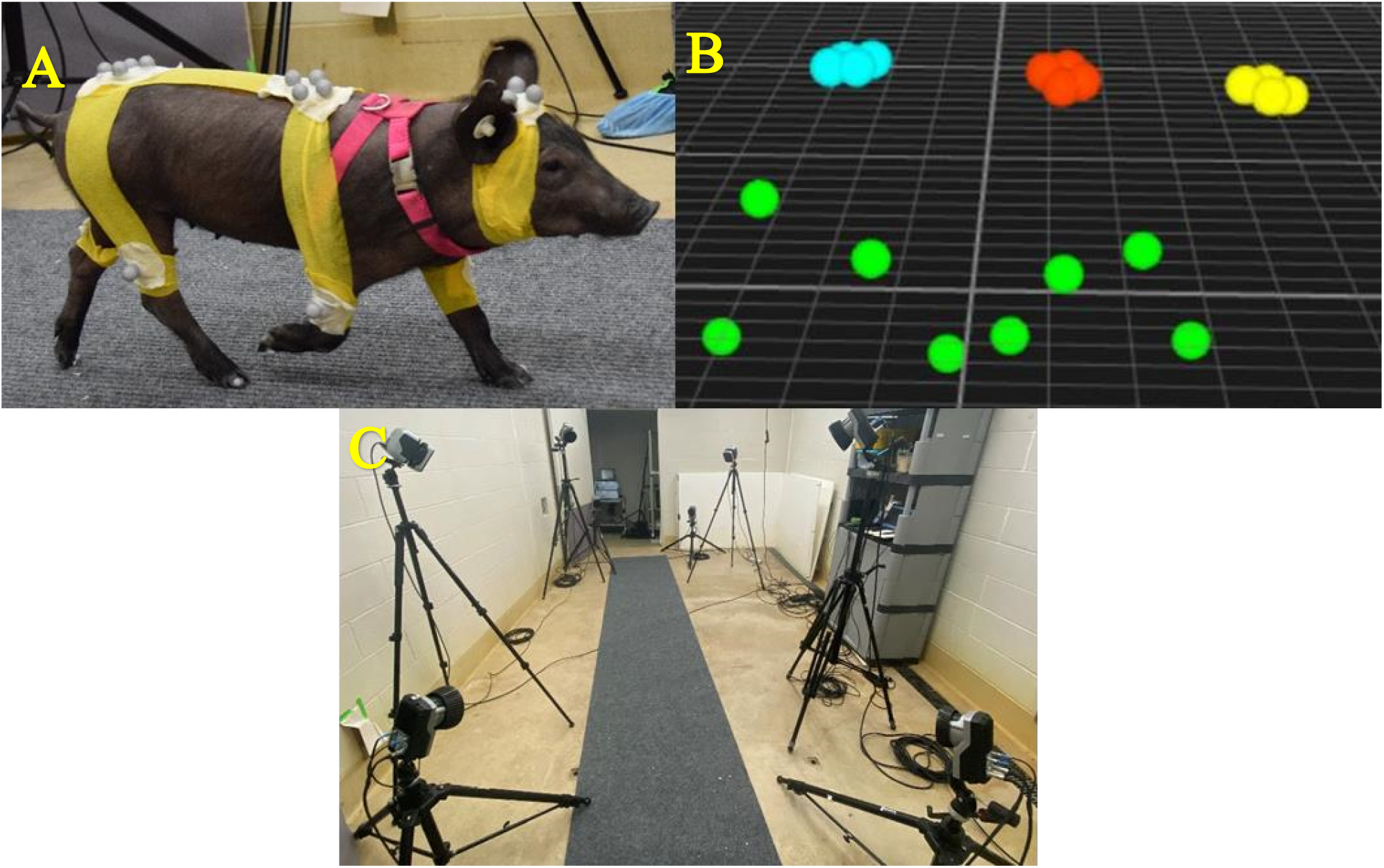
Motion capture setup. **A**. Retro-reflective markers were placed at different locations across the pig’s body. **B**. An example of rigid bodies captured by the system. **C**. The testing space.

### Hoof Height

A global coordinate system located on the ground provided a reference for the vertical height of each hoof marker. The hoof markers were placed laterally on the hoof towards as close to the base of the hoof as possible. In order to account for discrepancies in the placement of the hoof markers, the minimum height value for each hoof was set to the be the starting point to normalize, or the zero height, for that hoof marker. The reported vertical height of each hoof marker was then calculated using the measured hoof height minus the normalized minimum height of that hoof. The normalized hoof height was plotted against time for each successful pass of the pig.

### Swing and Stance Times + Gait Pattern of Steps

The pigs were able to walk at their own pace through the testing space, in order to reduce variables such as the speed of the pig, the cadence of the steps was determined for each pass. Only passes where the pigs were demonstrating a trotting gait pattern were used for comparison with one another. The trotting pattern was determined by the order of the steps relative to the other hooves in the normalized hoof height graphs. Using the hoof height graphs, the swing and stance times of each hoof were calculated. The swing time of a hoof was the time in which that hoof was off the ground. The stance time of a hoof was the time in which that hoof was placed on the ground. The swing time began at the last frame before the hoof began any vertical movement, and ended at the frame when the hoof ended any vertical movement. The stance time began as the swing time ended and the stance time ended when the hoof began to gain vertical height. The swing and stance times for multiple passes were averaged together to create the average swing and stance time for each pig.

### Rigid Body Rotations

Using methods defined in Grood and Suntay (33), the relative rotation angles for each marker pod were calculated. A local coordinate system was established on each marker pod using individual markers on each pod. When comparing the relative rotation of the head and the shoulders, the head acted as the fixed axis of rotation and the shoulders acted as the moving axis. When comparing the relative rotation of the shoulder and the hips, the shoulders acted the fixed axis of rotation, and the hips acted the moving axis. In order to gather the rotation data, a local coordinate system had to be defined for each marker pod. The local coordinate systems were defined with the x-axis from hips of pig toward head, y-axis from right side of pig to left side of pig, and z-axis goes vertically off the rigid body (from the ground upwards) (**Fig. 2A**).

**Figure 2.**
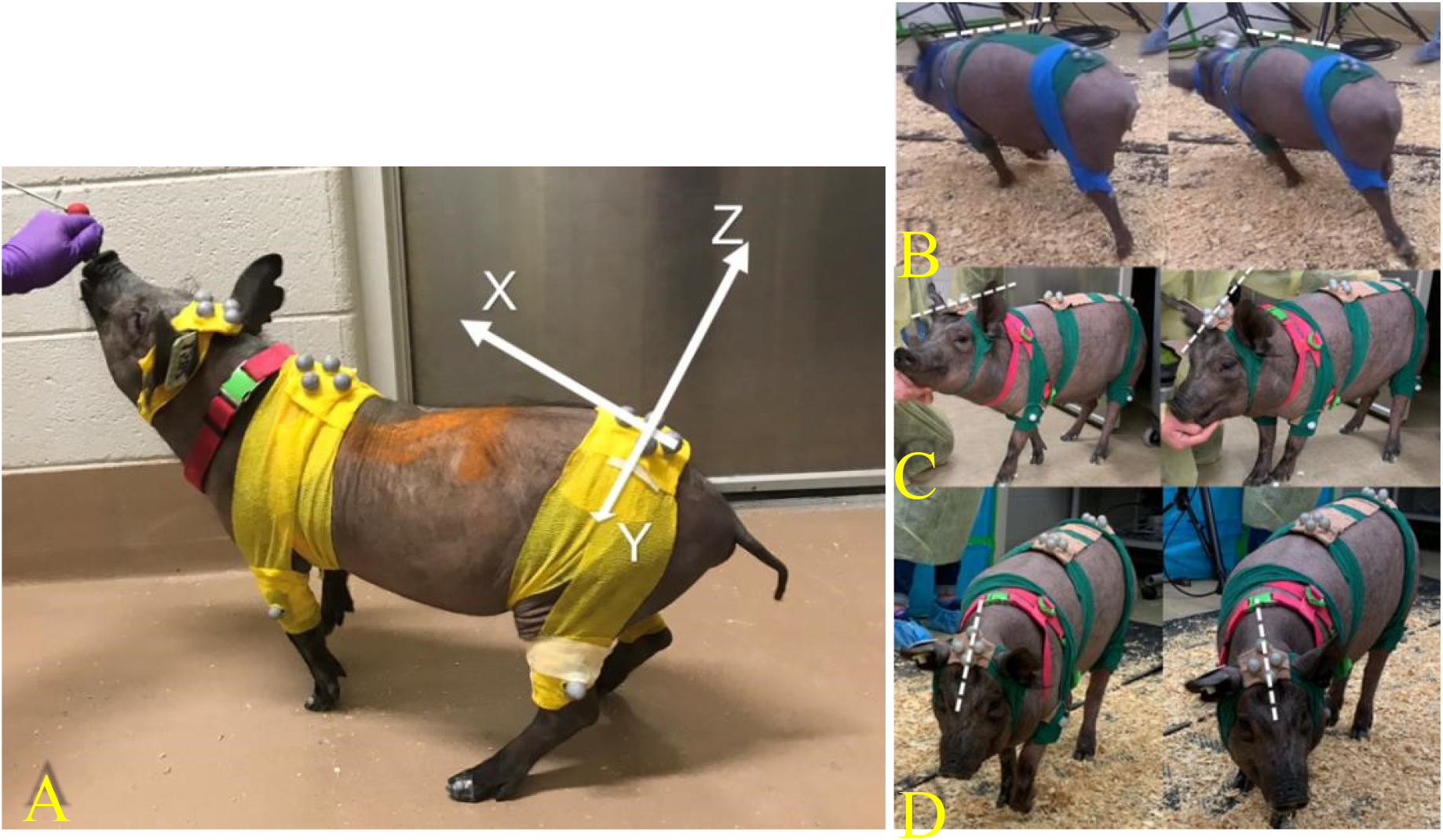
**A**. Local coordinate system placed onto the hip marker pod. **B**. Visual of the rotation about the x-axis. **C**. Visual of the rotation about the y-axis. **D**. Visual of the rotation about the z axis. The rotation around the x-axis was defined as roll, the rotation around the y-axis was defined as nodding, and the rotation around the z-axis was defined as side to side.

## Results

Measurements of body rotation in the Sham control pig, before and after the Sham procedures demonstrated that there was a maximum of 10 degrees of deviation in the nodding of the shoulders relative to the head, the rolling of the hips relative to the head, and the nodding of the hips relative to the head (**Table 1**).

**Table 1.** Average overall deviation angles of the shoulders relative to head and the hips relative to the head of a sham-control pig through multiple passes through the calibrated space.

|  |  | **Pre-procedure**<br>**(degrees)** | **After-Sham**<br>**procedure**<br>**(degrees)** |
| --- | --- | --- | --- |
| Shoulders to Head | Roll | 19.13 | 10.95 |
|  | Nodding | 29.23 | 39.56 |
|  | Side to Side | 14.33 | 10.09 |
| Hips to Head | Roll | 10.64 | 17.75 |
|  | Nodding | 40.10 | 48.74 |
|  | Side to Side | 10.60 | 11.81 |

The pigs that underwent the mTBI showed an increase in changes in angle deviations. As shown in **Table 2**, there were 16 degrees of deviation in the nodding of the shoulders relative to the head and 21 degrees of deviation in the nodding of the hips relative to the head.

**Table 2.** Average overall deviation angles of the shoulders relative to head and the hips relative to the head of a pig that did receive a TBI through multiple passes through the calibrated space.

|  |  | **Pre-procedure**<br>**(degrees)** | **Post-mTBI**<br>**procedure**<br>**(degrees)** |
| --- | --- | --- | --- |
| Shoulders to<br>Head | Roll | 15.43 | 9.66 |
|  | Nodding | 18.06 | 34.69 |
|  | Side to Side | 24.82 | 14.31 |
| Hips to Head | Roll | 19.54 | 7.66 |
|  | Nodding | 32.63 | 53.06 |
|  | Side to Side | 9.14 | 5.20 |

The pattern of the rotation angles changed between the pre- and post-TBI testing days (mTBI pig) is shown in **Figure 3**.

**Figure 3.**
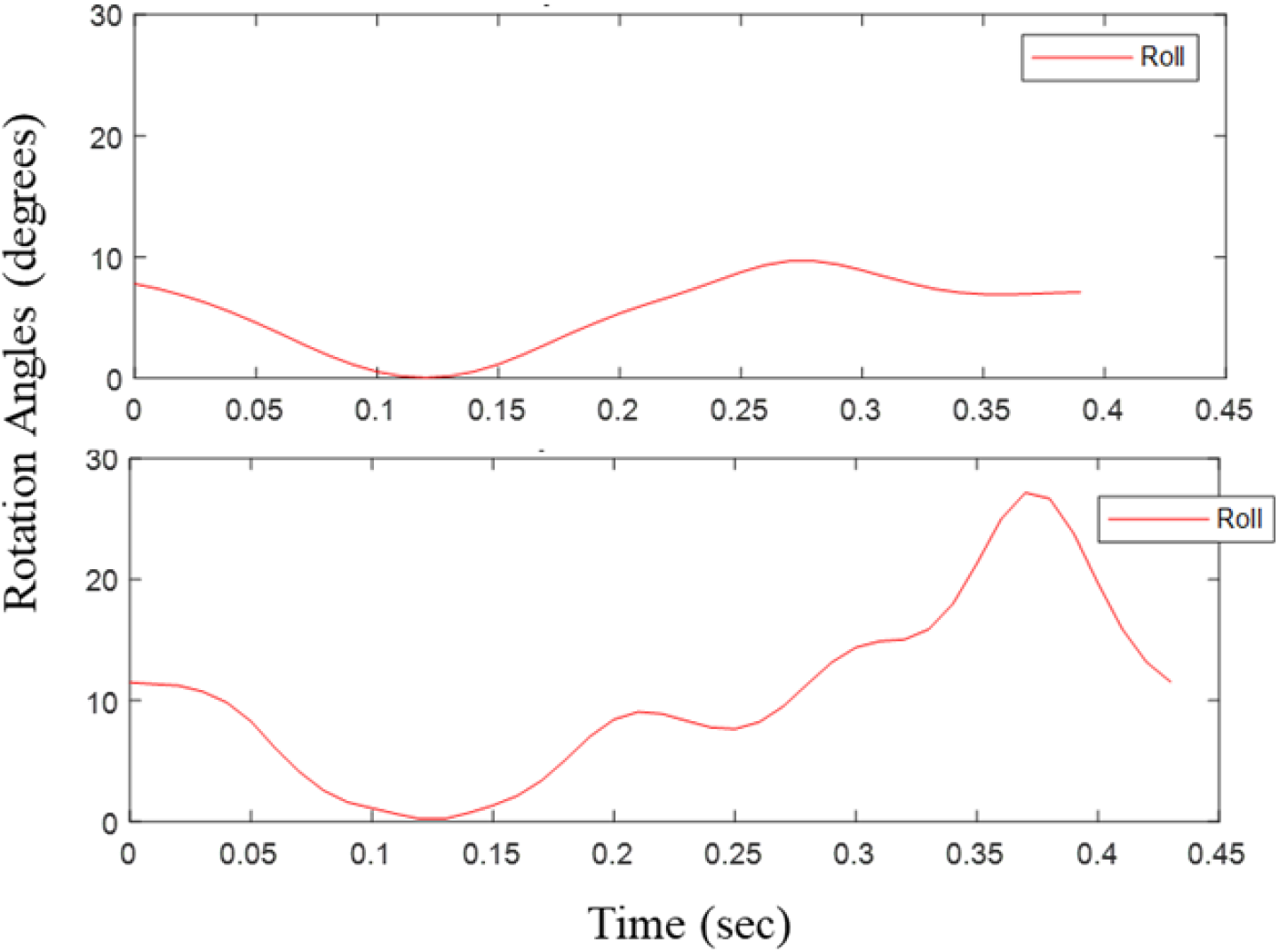
Rotation angles of one pass of a pig pre-TBI (top) and post-TBI (bottom) of the shoulders relative to the head.

**Figures 4** and **Figure 5** present the heights of each hoof, timing of steps, and comparison between the right and left side of the pig gait for one trial of the mTBI pig. The fore legs had an average swing time of 0.290±0.014 seconds, and the hind legs had an average swing time of 0.283±0.039 seconds.

**Figure 4.**
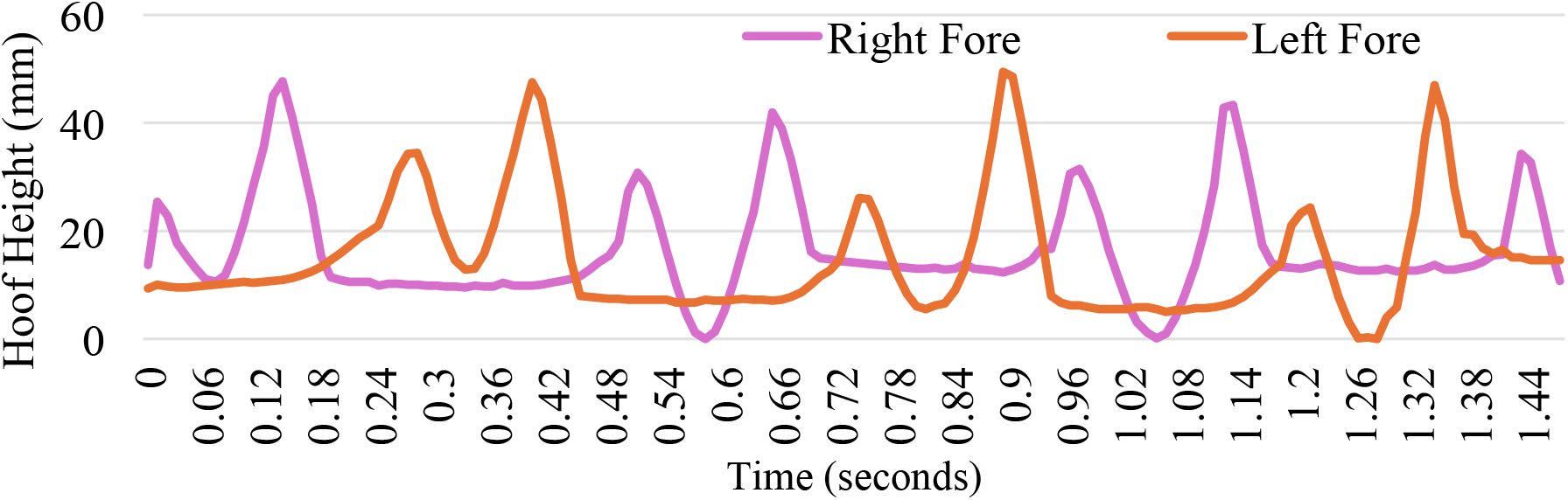
Fore-leg hoof height variation with time for one pig post-mTBI.

**Figure 5.**
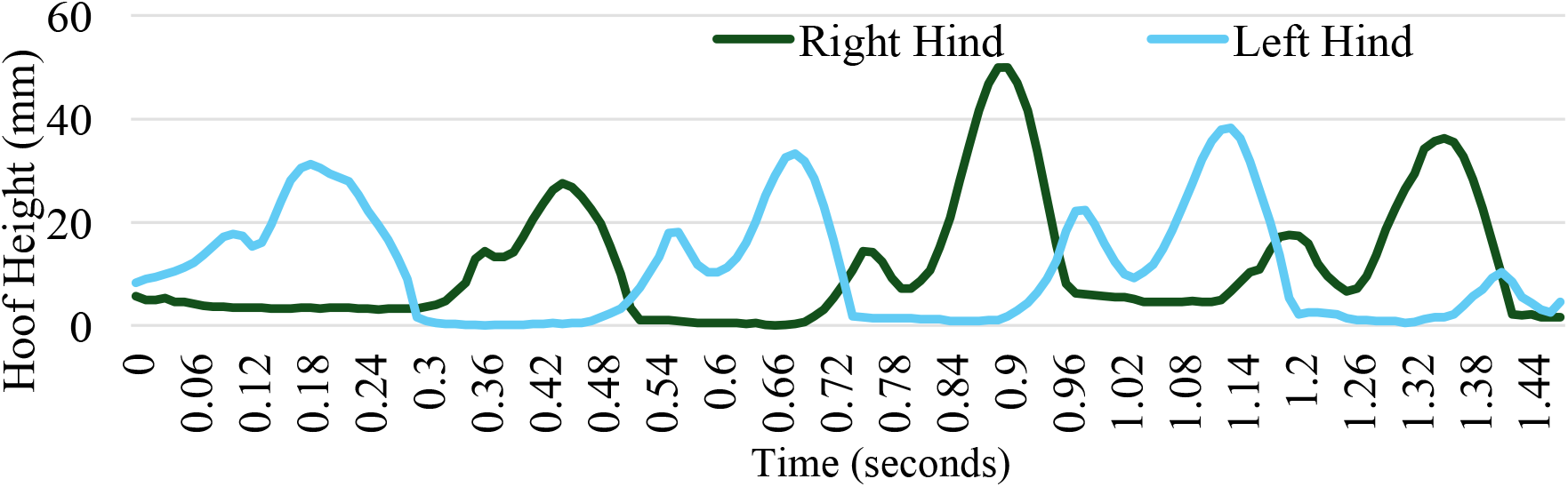
Hind-leg hoof height variation with time for one pig post-mTBI.

## Discussion

This study suggests that mTBI leads to changes in pig’s gait and body rotations. Shoulders relative to head means that the head was a fixed reference point, and the movement of shoulders were analyzed. The roll action was tilting side to side, the nodding action is moving up and down as the pig moves the head, and the side-to-side action is horizontal twisting such as turning head to the right or left. As shown in **Table 2**, the roll and side-to-side action decreased. The pig’s shoulders experienced less tilting and there was less lateral flexibility. Nodding movement was increased due to the shoulders moving up and down. These findings suggest potential instability in head control and a loss of fine motor control, similar to what is observed in human patients with TBI. The pigs may have been compensating for balance issues by restricting unnecessary side-to-side motion and increasing vertical, nodding, to help stabilize the head control.

Previous studies showed that TBI patients also experienced changes in center of pressure displacement with decreased velocity in medial/lateral planes (34). In agreement with these studies, the mTBI pig demonstrated a decreased side to side movement due to center of pressure instability. The Roll movement is leaning left or right, nodding is up and down movement as the pig’s hips were moving vertically, and the side-to-side motion is rotation of hips by twisting movement. This movement decreased as shown in **Table 2** which could be due to reduced lateral speed and flexibility. The side-to-side decrease is likely due to stiffness and balance issues. The nodding increased observed in the pig aligns with the observations of unsteadiness in human TBI individuals (35). Individuals who suffered TBI also expend more energy to maintain balance through compensatory mechanisms to achieve stability (11). Athletes also reported altered gait patterns, gait velocity, and poorer gait balance control during walking post TBI (36).

In our study, the observations in changes in body rotations infer that TBI affected the pig’s gait. There was a decreased side-to-side movement usually means controlled walking and shorter steps and slower movement. The pigs were moving more cautiously which could infer with increased stance duration by keeping their feet on the ground longer for stability. A previous study assessing gait impairment following TBI injury in a mouse model showed similar findings; The mice with TBI had increased stance and swing duration and decreased stride length and gait velocity (37).

There were many gait changes observed in the mTBI pig. One main reason could be due neuromuscular impairments. It is well documented that TBI often leads to deficits in function and biomechanical patterns. Aberrant kinematics, abnormal movement patterns, and balance impairments are also altered due to the impact on the brain (38). The vestibular system is also altered through the impact. Impairments related to TBI include vertigo, dizziness, balance, gait disorders, blurry vision, and others in humans (39). The pigs could have had an impairment to the vestibular balance and tried to compensate through increased nodding to adjust their balance.

While this study provided insights into changes in gait associated with mTBI, there are several limitations that should be considered. In the study, only two pigs were tested: one for control and one for post-TBI. Having a larger sample size will be required to determine the variability of movement in each group and statistical analysis of the results.

Even though a pig is a good translational model, there are still many differences between pigs and humans. Pigs have four legs, two hind and fore legs, whereas humans just have two legs. Humans gait impairments post TBI involve speed, balance, step length, and many other factors whereas pigs may compensate differently or show different symptoms due to their four legs. There are also different environmental and emotional disturbances that can affect TBI recovery and symptoms for humans that pigs may not experience (40).

Because of the complexity of evaluating the gait of pigs, the majority of the published research has focused on hoof contact patterns. There is limited TBI related gait research, particularly with the inclusion on relative movements between body segments. This work demonstrates the ability to gather hoof height, timing of each leg during gait and relative motions between body segments. All of these measures have the potential to provide insights to changes in movements pre- and post-TBI. Determining not only magnitude of movements, but timing of these movements will add to our understanding of functional changes in gait patterns. This is the initial step for a full evaluation of multiple pigs pre- and post-TBI injury.

This study successfully developed methods for the gathering of natural gait data from Yucatan minipigs after TBI and shows how mTBI may induce imbalance in this model, similar to what is seen in the clinic. The findings highlight the utility of the Yucatan minipig as a relevant animal model for studying mild traumatic brain injury and its effects on postural control and gait. Through detailed kinematic and gait analysis, we were able to identify specific alterations in movement patterns, such as decreased shoulder flexibility and compensatory head stabilization, which parallel the symptoms observed in human patients with mTBI. These results not only enhance our understanding of the physiological changes that occur following mild TBI but also offer a valuable framework for assessing balance impairments in future therapeutic interventions. Overall, this study contributes to the growing body of research aimed at improving the diagnosis and treatment of balance-related disorders in both animal models and human patients.

## Acknowledgments

N/A

## Competing Interests

The authors have declared that no competing interests exist.

## Notes

### Competing Interest Statement

The authors have declared no competing interest.

## Bibliography

1. A. Capizzi, J. Woo, M. Verduzco-Gutierrez, Traumatic Brain Injury: An Overview of Epidemiology, Pathophysiology, and Medical Management. Med Clin North Am 104, 213–238 (2020).

2. M. J. Haydel, L. J. Weisbrod, W. Saeed, “Pediatric Head Trauma” in StatPearls. (Treasure Island (FL) ineligible companies. 2025).

3. K. O. Yeates et al., Short- and long-term social outcomes following pediatric traumatic brain injury. J Int Neuropsychol Soc 10, 412–426 (2004).

4. B. Wechsler, H. Kim, P. R. Gallagher, C. DiScala, M. G. Stineman, Functional status after childhood traumatic brain injury. J Trauma 58, 940–949; discussion 950 (2005).

5. A. McKinlay, R. C. Grace, L. J. Horwood, D. M. Fergusson, M. R. MacFarlane, Long-term behavioural outcomes of pre-school mild traumatic brain injury. Child Care Health Dev 36, 22–30 (2010).

6. D. M. Ransom et al., Academic effects of concussion in children and adolescents. Pediatrics 135, 1043–1050 (2015).

7. L. Ewing-Cobbs et al., Modeling of longitudinal academic achievement scores after pediatric traumatic brain injury. Dev Neuropsychol 25, 107–133 (2004).

8. G. L. Iverson, P. M. Kelshaw, N. E. Cook, S. V. Caswell, Middle School Children With Attention-Deficit/Hyperactivity Disorder Have a Greater Concussion History. Clin J Sport Med 31, 438–441 (2021).

9. K. Sambasivan, L. Grilli, I. Gagnon, Balance and mobility in clinically recovered children and adolescents after a mild traumatic brain injury. J Pediatr Rehabil Med 8, 335–344 (2015).

10. D. R. Howell, L. R. Osternig, L. S. Chou, Dual-task effect on gait balance control in adolescents with concussion. Arch Phys Med Rehabil 94, 1513–1520 (2013).

11. A. Al-Husseini et al., Long-term postural control in elite athletes following mild traumatic brain injury. Front Neurol 13, 906594 (2022).

12. K. R. Campbell et al., Assessment of balance in people with mild traumatic brain injury using a balance systems model approach. Gait Posture 100, 107–113 (2023).

13. K. R. Campbell et al., Exploring persistent complaints of imbalance after mTBI: Oculomotor, peripheral vestibular and central sensory integration function. J Vestib Res 31, 519–530 (2021).

14. S. S. Shin et al., Transcranial magnetic stimulation and environmental enrichment enhances cortical excitability and functional outcomes after traumatic brain injury. Brain Stimul 11, 1306–1313 (2018).

15. N. Li et al., Evidence for impaired plasticity after traumatic brain injury in the developing brain. J Neurotrauma 31, 395–403 (2014).

16. H. Lu et al., Transcranial magnetic stimulation facilitates neurorehabilitation after pediatric traumatic brain injury. Sci Rep 5, 14769 (2015).

17. S. R. Shultz et al., The potential for animal models to provide insight into mild traumatic brain injury: Translational challenges and strategies. Neurosci Biobehav Rev 76, 396–414 (2017).

18. C. Vonder Haar et al., Repetitive closed-head impact model of engineered rotational acceleration (CHIMERA) injury in rats increases impulsivity, decreases dopaminergic innervation in the olfactory tubercle and generates white matter inflammation, tau phosphorylation and degeneration. Exp Neurol 317, 87–99 (2019).

19. A. H. Netzley, G. Pelled, The Pig as a Translational Animal Model for Biobehavioral and Neurotrauma Research. Biomedicines 11 (2023).

20. A. H. Netzley et al., Multimodal characterization of Yucatan minipig behavior and physiology through maturation. Sci Rep 11, 22688 (2021).

21. S. Islam et al., Volumetric and Diffusion Tensor Imaging biomarkers indicating long lasting post-concussion abnormalities in a youth pig model of mild Traumatic Brain Injury. BioRxiv (2024).

22. D. K. Cullen et al., A Porcine Model of Traumatic Brain Injury via Head Rotational Acceleration. Methods Mol Biol 1462, 289–324 (2016).

23. A. R. Mayer et al., Survival Rates and Biomarkers in a Large Animal Model of Traumatic Brain Injury Combined With Two Different Levels of Blood Loss. Shock 55, 554–562 (2021).

24. J. A. Wolf et al., Concussion Induces Hippocampal Circuitry Disruption in Swine. J Neurotrauma 34, 2303–2314 (2017).

25. E. W. Baker et al., Scaled traumatic brain injury results in unique metabolomic signatures between gray matter, white matter, and serum in a piglet model. PLoS One 13, e0206481 (2018).

26. E. W. Baker et al., Controlled Cortical Impact Severity Results in Graded Cellular, Tissue, and Functional Responses in a Piglet Traumatic Brain Injury Model. J Neurotrauma 36, 61–73 (2019).

27. G. Simchick et al., Detecting functional connectivity disruptions in a translational pediatric traumatic brain injury porcine model using resting-state and task-based fMRI. Sci Rep 11, 12406 (2021).

28. H. Wang et al., Identification of predictive MRI and functional biomarkers in a pediatric piglet traumatic brain injury model. Neural Regen Res 16, 338–344 (2021).

29. K. J. Duberstein et al., Gait analysis in a pre- and post-ischemic stroke biomedical pig model. Physiol Behav 125, 8–16 (2014).

30. B. B. Reynolds et al., Quantifying Head Impacts in Collegiate Lacrosse. Am J Sports Med 44, 2947–2956 (2016).

31. S. B. Sandmo, A. S. McIntosh, T. E. Andersen, I. K. Koerte, R. Bahr, Evaluation of an In-Ear Sensor for Quantifying Head Impacts in Youth Soccer. Am J Sports Med 47, 974–981 (2019).

32. J. Kerwin et al., Sulcal Cavitation in Linear Head Acceleration: Possible Correlation With Chronic Traumatic Encephalopathy. Front Neurol 13, 832370 (2022).

33. E. S. Grood, W. J. Suntay, A joint coordinate system for the clinical description of three-dimensional motions: application to the knee. J Biomech Eng 105, 136–144 (1983).

34. M. Bonanno et al., Gait Analysis in Neurorehabilitation: From Research to Clinical Practice. Bioengineering (Basel) 10 (2023).

35. J. Row et al., Balance Assessment in Traumatic Brain Injury: A Comparison of the Sensory Organization and Limits of Stability Tests. J Neurotrauma 36, 2435–2442 (2019).

36. D. R. Howell, R. C. Lynall, T. A. Buckley, D. C. Herman, Neuromuscular Control Deficits and the Risk of Subsequent Injury after a Concussion: A Scoping Review. Sports Med 48, 1097–1115 (2018).

37. M. Neumann et al., Assessing gait impairment following experimental traumatic brain injury in mice. J Neurosci Methods 176, 34–44 (2009).

38. R. C. Lynall et al., Investigating post-mild traumatic brain injury neuromuscular function and musculoskeletal injury risk: A protocol for a prospective, observational, case-controlled study in service members and active individuals. BMJ Open 13, e069404 (2023).

39. E. Galeno, E. Pullano, F. Mourad, G. Galeoto, F. Frontani, Effectiveness of Vestibular Rehabilitation after Concussion: A Systematic Review of Randomised Controlled Trial. Healthcare (Basel) 11 (2022).

40. Q. Zhao, J. Zhang, H. Li, H. Li, F. Xie, Models of traumatic brain injury-highlights and drawbacks. Front Neurol 14, 1151660 (2023).

